# A Multistage Sequencing Strategy Pinpoints Many Novel and Candidate Disease Alleles for Orphan Disease Emery-Dreifuss Muscular Dystrophy and Supports Gene Misregulation as its Pathomechanism

**DOI:** 10.1101/705780

**Authors:** Peter Meinke, Alastair R. W. Kerr, Rafal Czapiewski, Jose I. de las Heras, Elizabeth Harris, Heike Kölbel, Francesco Muntoni, Ulrike Schara, Volker Straub, Benedikt Schoser, Manfred Wehnert, Eric C. Schirmer

**Affiliations:** Wellcome Centre for Cell Biology, University of Edinburgh, Edinburgh, UK; Friedrich Baur Institute at the Department of Neurology, University Hospital, LMU Munich, Germany; John Walton Muscular Dystrophy Research Centre, Newcastle University and Newcastle Hospitals NHS Foundation Trust, Newcastle upon Tyne, UK; Department of Pediatric Neurology, Developmental Neurology and Social Pediatrics, University of Essen, Germany; Dubowitz Neuromuscular Centre, University College London Great Ormond Street Institute of Child Health, London, UK; and 1 NIHR Great Ormond Street Hospital Biomedical Research Centre, Great Ormond Street Institute of Child Health, University College London, & Great Ormond Street Hospital Trust, London, UK; Institute of Human Genetics, University of Greifswald (retired), Greifswald, Germany

**Keywords:** Emery-Dreifuss muscular dystrophy, nuclear envelope, nuclear envelope transmembrane protein, primer library, orphan disease

## Abstract

Limitations of genome-wide approaches for genetically-heterogenous orphan diseases led us to develop a new approach to identify novel Emery-Dreifuss muscular dystrophy (EDMD) candidate genes. We generated a primer library to genes: (I) linked to EDMD, (II) mutated in related muscular dystrophies, (III) highlighted from limited exome sequencing, (IV) encoding muscle-specific nuclear membrane proteins. Sequencing 56 unlinked EDMD patients yielded confirmed or strong candidate alleles from all categories, accounting for most remaining unlinked patients. Known functions of newly-linked genes argue the EDMD pathomechanism is from altered gene regulation and mechanotransduction through connectivity of candidates from the nuclear envelope to the plasma membrane.

---

Emery-Dreifuss muscular dystrophy (EDMD) is a rare neuromuscular disorder affecting ∼0.3-0.4 in 100,000 people^1, 2^. EDMD patients present typically in childhood with early contractures of elbows and Achilles’ tendons, progressive wasting of lower leg and upper arm muscles, and later development of cardiac conduction defects and, in a proportion of cases, dilated cardiomyopathy^3^. Features vary considerably in clinical presentation, leading to the usage ‘Emery-Dreifuss-like syndromes’^4, 5^: patients from the same pedigree can show remarkable phenotypic variation^6–8^. Consistent with this variation, EDMD is also genetically variable: ∼half of Emery–Dreifuss-like syndrome cases are linked to mutations in genes encoding 6 nuclear envelope proteins (emerin, lamin A, nesprin 1, nesprin 2, SUN1 and FHL1^8–12^). Variants in desmin and nuclear envelope proteins Tmem43 and SUN2 have been reported to modify the EDMD phenotype^11, 13, 14^. Roughly half of clinically diagnosed patients remain unlinked^15, 16^.

The strong nuclear envelope link raised the possibility that remaining unlinked patients might also have mutations in nuclear envelope proteins. The nuclear envelope is linked to >30 inherited diseases and syndromes^17^, each with distinct tissue-specific pathologies: for example different lamin A mutations cause muscular dystrophies, neuropathy, lipodystrophy, and multisystemic disorders^18^. How these widely expressed nuclear envelope proteins yield tissue-specific pathologies remains unresolved, but one hypothesis is that tissue-specific nuclear envelope partners mediate the tissue-specificity of effects^18, 19^.

We previously identified several muscle-specific nuclear envelope transmembrane proteins (NETs)^20^. Of the previously linked proteins emerin, nesprin 1, nesprin 2, SUN1, SUN2, and Tmem43 are all NETs, but these are widely expressed. Several of the muscle-specific NETs identified could contribute muscle specificity to either of the two principly hypothesized EDMD pathomechanisms: mechanical instability and disruption of gene expression^21^. NETs Tmem214 and KLHL31 track with microtubules on the nuclear surface^20^ while NET5/Samp1 contributes actin and centrosome interactions^22, 23^. NETs Tmem38A, WFS1, NET39/PLPP7 and, again, Tmem214 and NET5/Samp1 all affect 3D gene positioning and with corresponding effects on expression^24, 25^. Many of the genes under muscle-specific NET regulation are recruited to the nuclear periphery to be more tightly shut down during myogenesis and encode proteins that are antagonistic to myogenesis or are from alternative differentiation pathways such as adipocytes. Knockdown of the muscle-specific NETs results in these genes being de-repressed, suggesting a possible gene misregulation mechanism to disease pathology. The potential of gene mispositioning contributing to disease is further underscored by knockdown of Tmem38A, WFS1, and NET39/PLPP7 blocking myotube fusion^24^. Functional overlap of these muscle-specific NETs supports the possibility of their working in a common pathway towards EDMD pathophysiology, making them good candidates for mediators of EDMD muscle pathology at the same time as being novel candidates for causative EDMD alleles.

Therefore, we elected to sequence the genes encoding these muscle-specific NETs in unlinked EDMD patients using a primer library. However, for greater surety, we expanded this primer library to also re-check previously linked genes with complete gene sequencing for possible promoter mutations and to test for mutations in genes linked to related muscular dystrophies. Finally, to also search for candidate alleles in a completely unbiased manner, we performed exome sequencing in families for which material from enough members was available for linkage analysis and added these candidates also to the primer library.

## Results

### SEQUENCING EDMD FAMILIES

Whole exome sequencing was performed in 12 EDMD patients and 12 unaffected individuals from 5 families with large enough pedigrees for linkage analysis (Fig. 1), finding over 250,000 variants compared to the reference sequence. Variants were filtered using criteria: (a) phenotype co-segregation and modes of inheritance for each family; (b) selecting for SNP frequencies <1%, and filtering for <0.05%; (c) affecting coding sequence; (d) function/tissue-expression of the encoded protein *e.g.* >2-fold higher expression in muscle compared to other tissues. Filtering yielded 213 candidate genes for families 2-5 (Supplemental Table S1).

**Figure 1.**
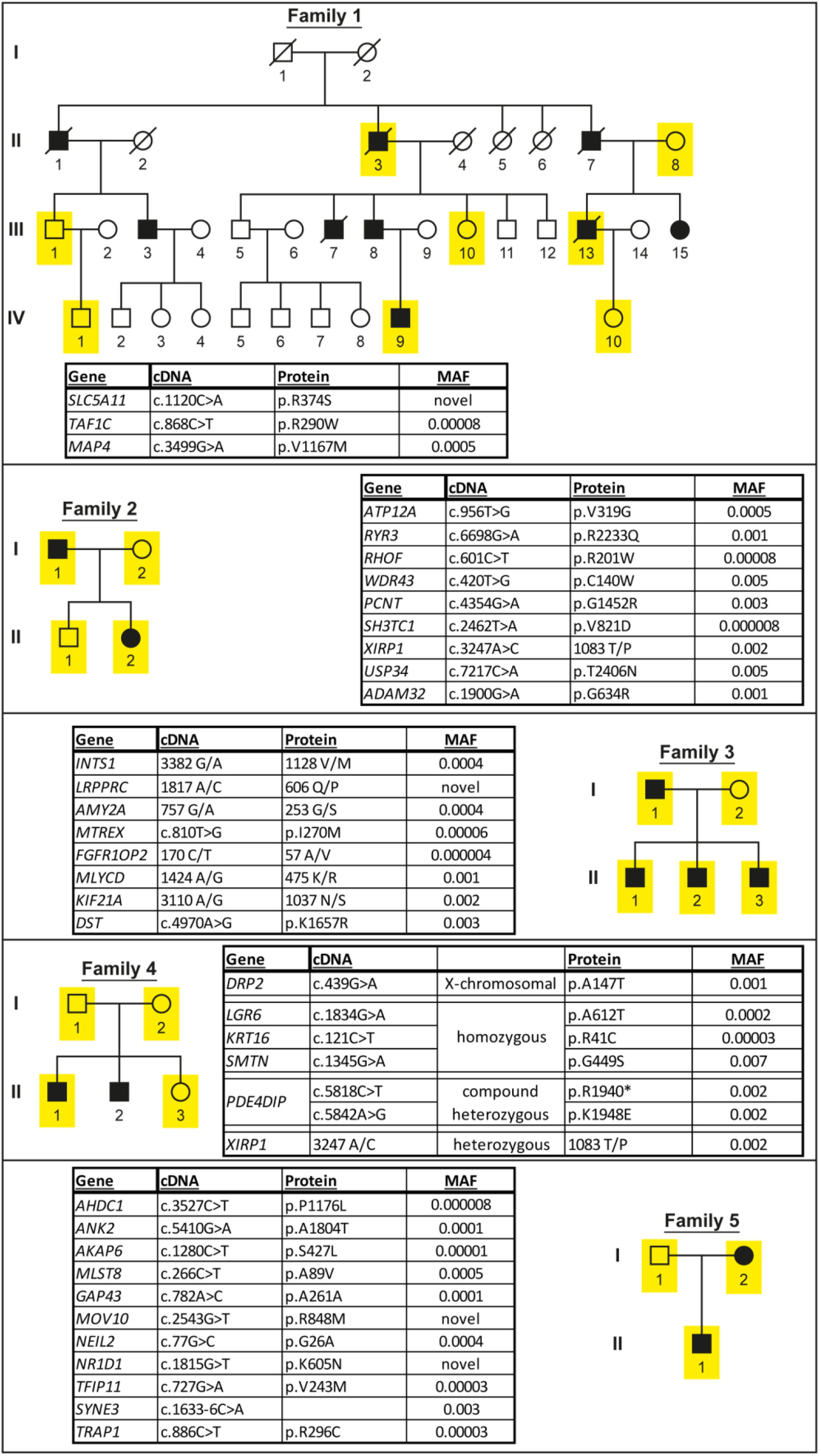
Pedigrees of the 5 families used for the initial exome sequencing with top candidate mutations listed in the adjacent boxes (MAF = minor allele frequency). Sequenced individuals: yellow; males: square; females: circles; patients: filled black.

Family 1 yielded no convincing candidates. As this family had the largest pedigree, we postulated that an unaffected individual was a carrier who had not yet presented or had a distinct sporadic form of disease. Dropping younger individuals who may have not yet presented clinically failed to yield candidates; therefore, genome and transcriptome sequencing was performed in the index patient, resulting in 33 additional candidates (Supplemental Tables S2 and S3). The combined exome, genome and RNA sequencing yielded a total of 252 candidates for the five families.

## PRIMER LIBRARY SEQUENCING

A primer library was generated containing (I) the 8 previously-linked EDMD gene ORFs plus the whole genes for *LMNA* and *EMD* (that together account for ∼40% of linked alleles), (II) 25 genes from similar muscular dystrophies, (III) the 252 exome sequencing candidates, and (IV) 16 functional candidates, mostly muscle-specific nuclear envelope proteins (Fig. 2A; Supplemental Table S4). Sequencing was performed on 56 additional unlinked clinically diagnosed EDMD patients unrelated to each other, obtaining on average 3,427,092 reads per patient. The data were analyzed for genes carrying mutations that changed the coding sequence (nonsense, missense, splice sites) with expected altered protein function (e.g. non-conservative substitutions) and SNP frequencies <0.05% (Supplemental Table S5).

**Figure 2.**
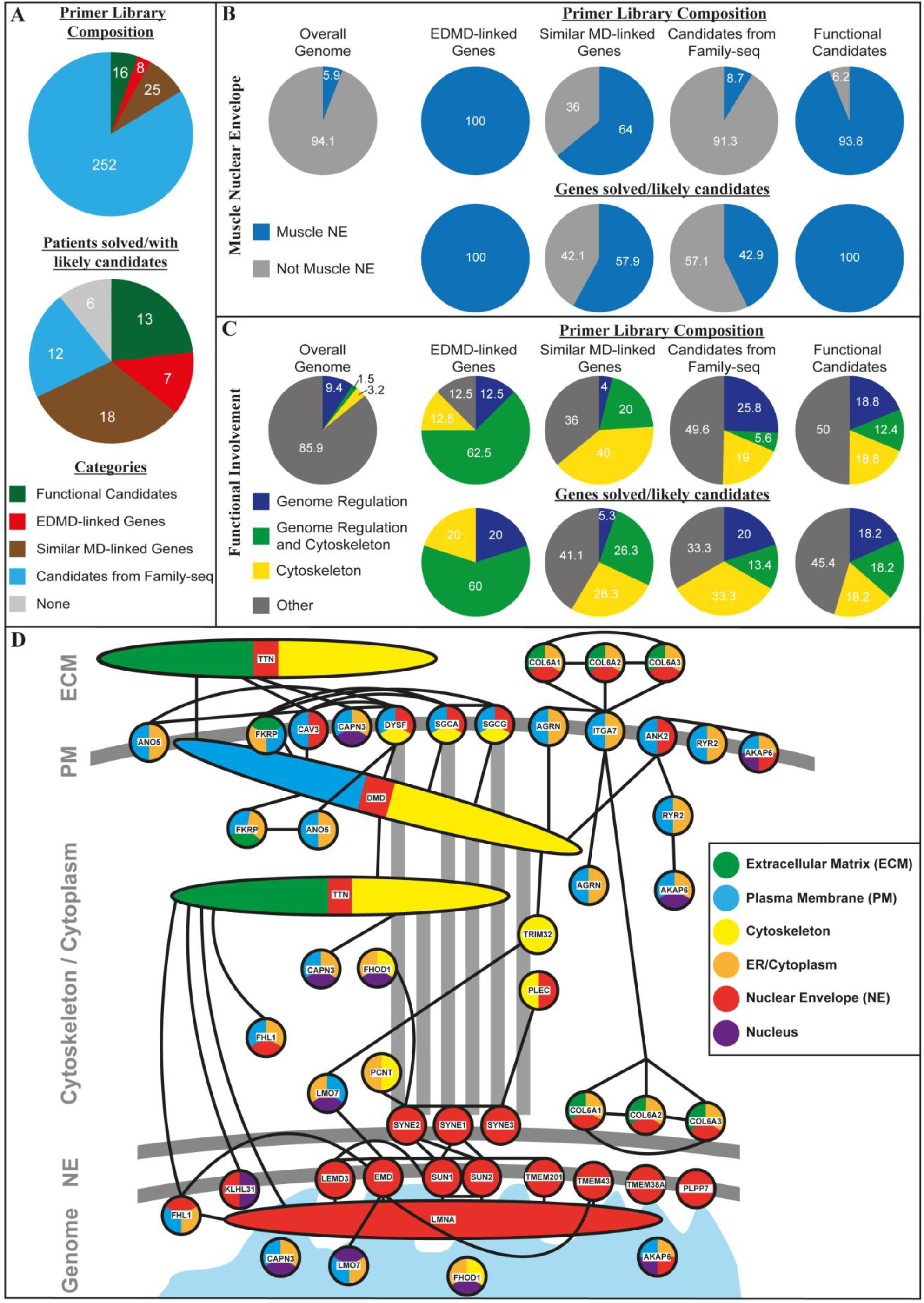
Primer library composition and gene ontology (GO) functions/localizations of all candidate genes from the four categories contributing to the primer library construction and for the top candidates identified after primer library sequencing. **(A)** Composition of the primer library with number of genes from each of the four categories used in its construction (upper panel) and number of patients solved/with likely candidates from the different categories after primer library sequencing (lower panel). **(B)** Presence in muscle nuclear envelopes for the starting library in comparison to the overall genome (upper panel) and of the remaining candidate genes after primer library sequencing (lower panel) in percent (based on GO-localization terms and/or experimental evidence from appearance in nuclear envelope proteomics datasets^20, 28^). **(C)** GO-terms for genome organization, cytoskeleton, and genome organization and cytoskeleton combined functions involvement for the starting library in comparison to the overall genome (upper panel) and of the remaining candidate genes after primer library sequencing (lower panel) in percent, showing an enrichment for the combined category in the top candidate alleles. **(D)** Interactive network of remaining candidate genes after library sequencing based on STRING (Search Tool for the Retrieval of Interacting Genes/Proteins, https://string-db.org/) interactions (high confidence) showing that most candidates are linked to other candidates and that these form connections from the nuclear envelope to the plasma membrane. These connections are consistent with possible mechanotransduction from the extracellular region to the nuclear envelope being the core disrupted function in EDMD. Different described localizations of proteins are displayed by color-coding (based on GO-terms and/or experimental evidence through identification in muscle nuclear envelope proteomics datasets^20^).

Candidate mutations were found in all four categories. Of category I previous EDMD-linked genes, *LMNA* had mutations in three patients that were missed in standard diagnostics (p.R41H, p.R249Q, p.G535fs*; Table 1). These mutations were determined as causative based on similarity to previously linked *LMNA* mutations. Previously EDMD-associated genes *SYNE1*, *SYNE2*, *SUN1* and *TMEM43* also had mutations; however, minor allele frequencies and their combination with other mutations made them unlikely as causative alleles excepting *SYNE1*. Modifying effects, nonetheless, cannot be excluded. No mutations were found in *LMNA* or *EMD* non-coding regions.

**Table 1:**
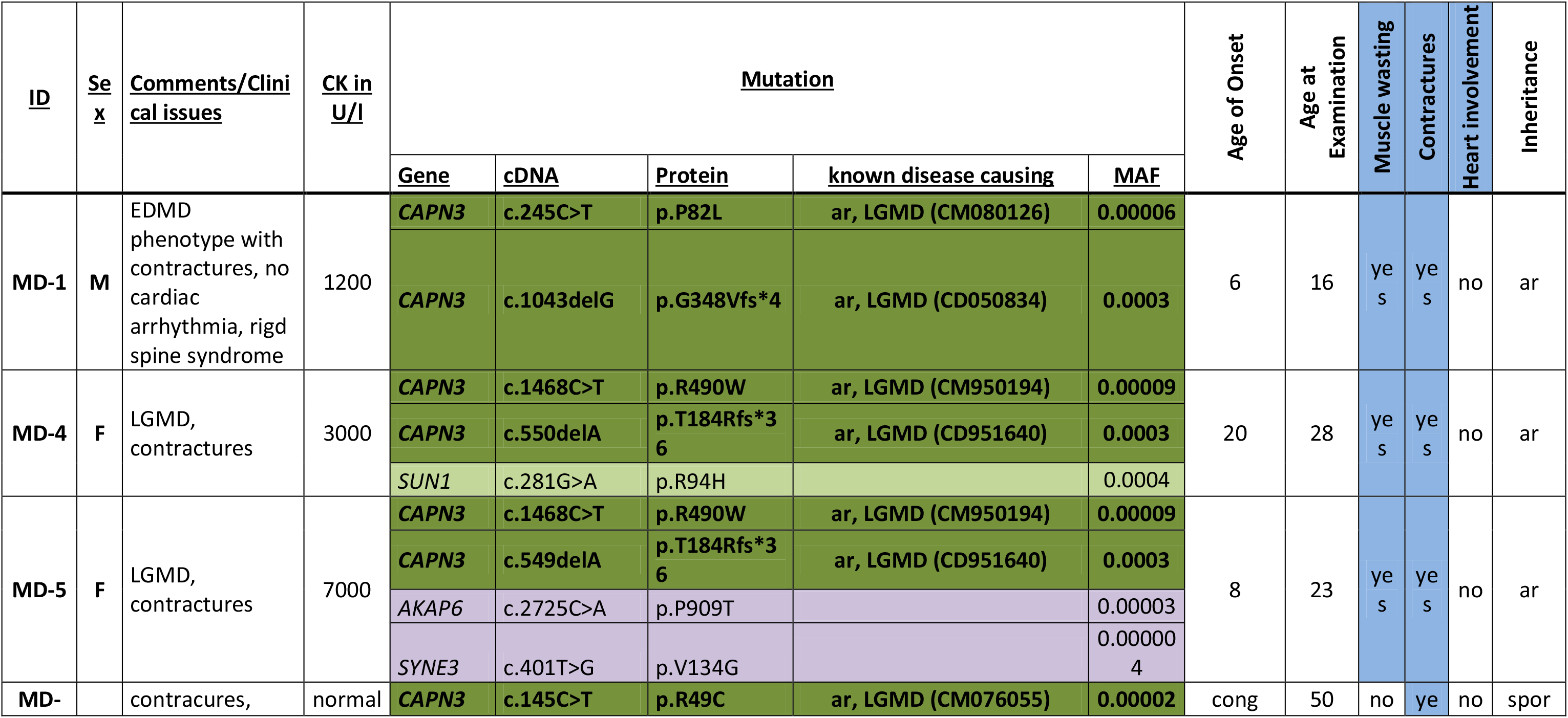
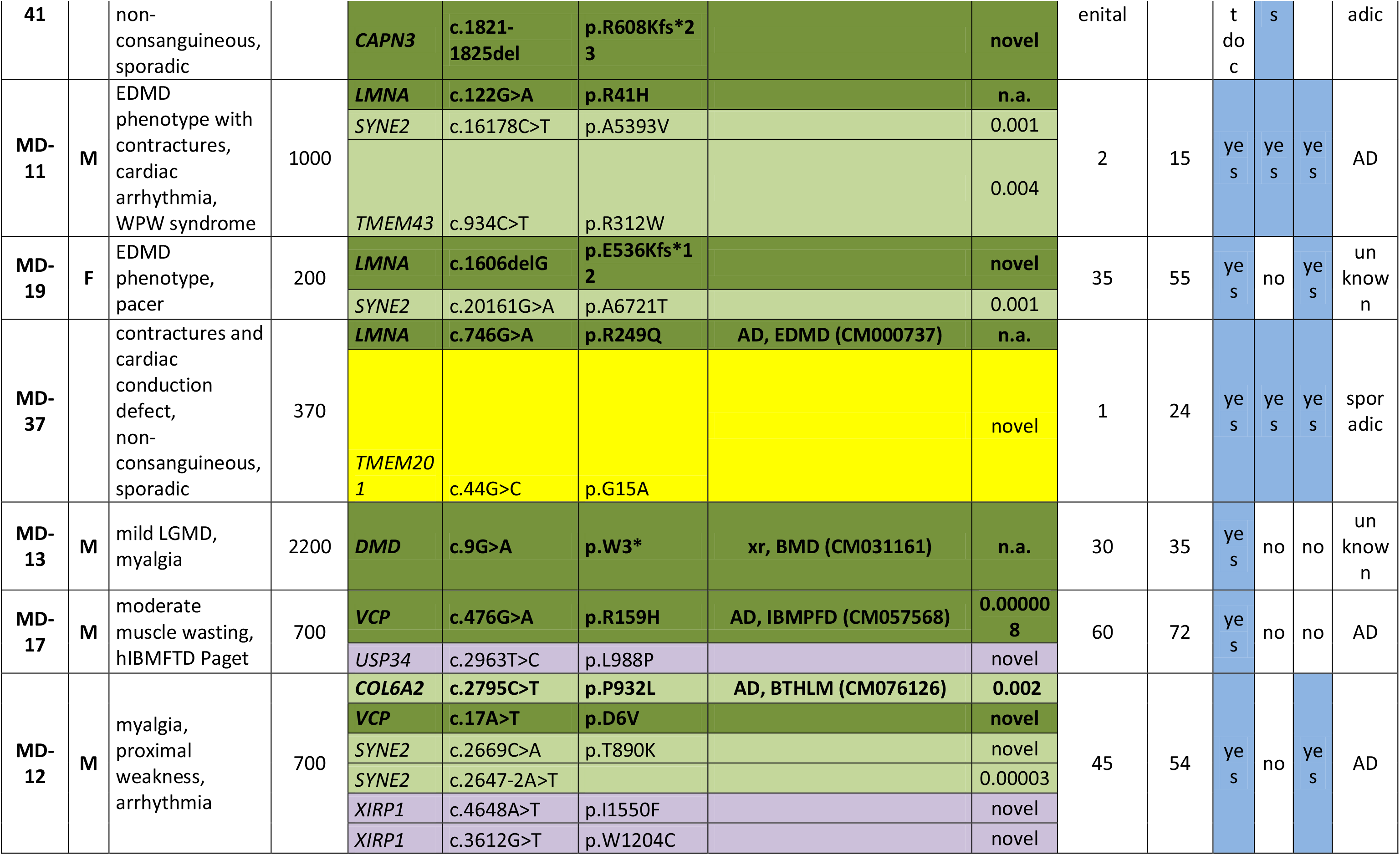
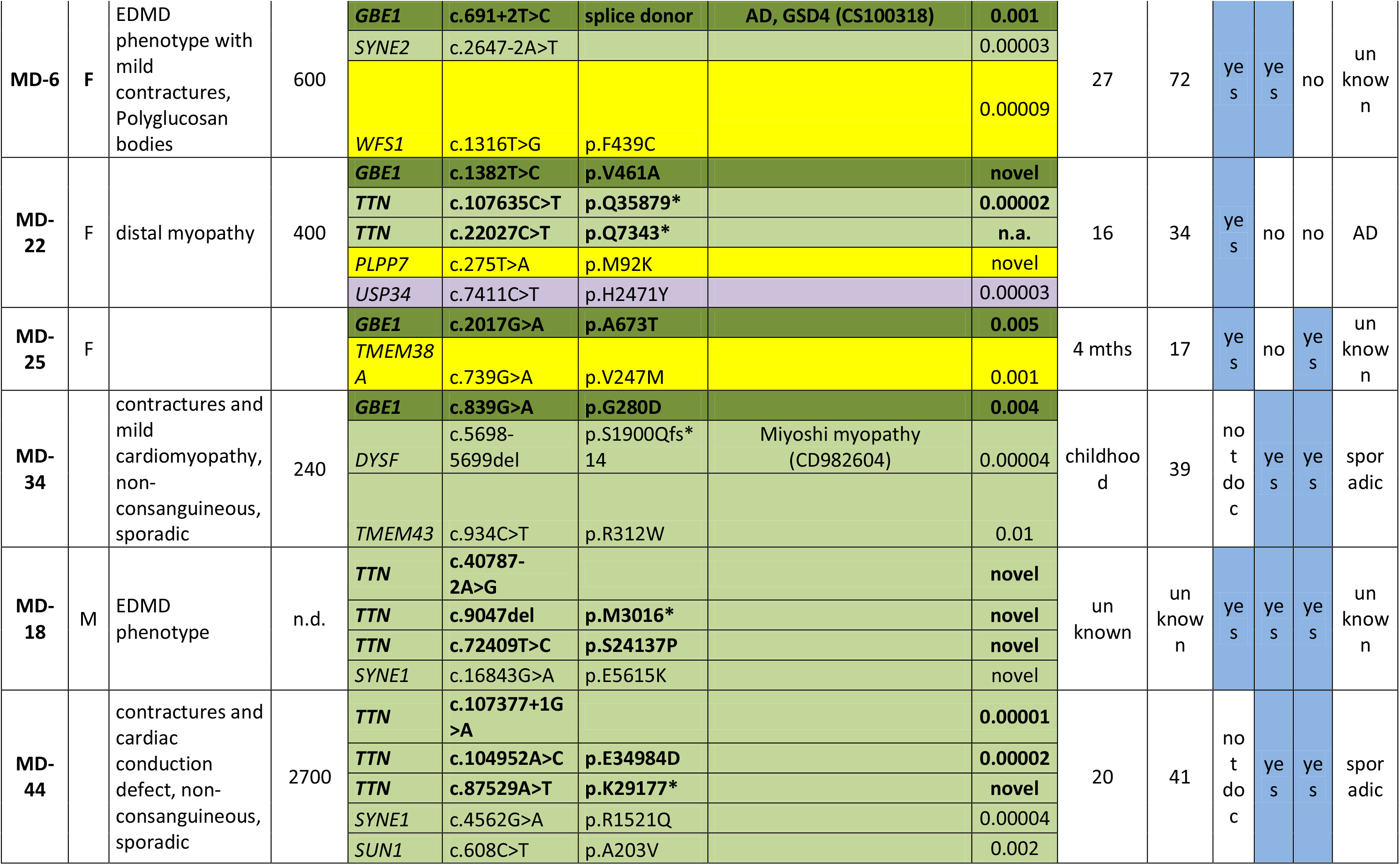
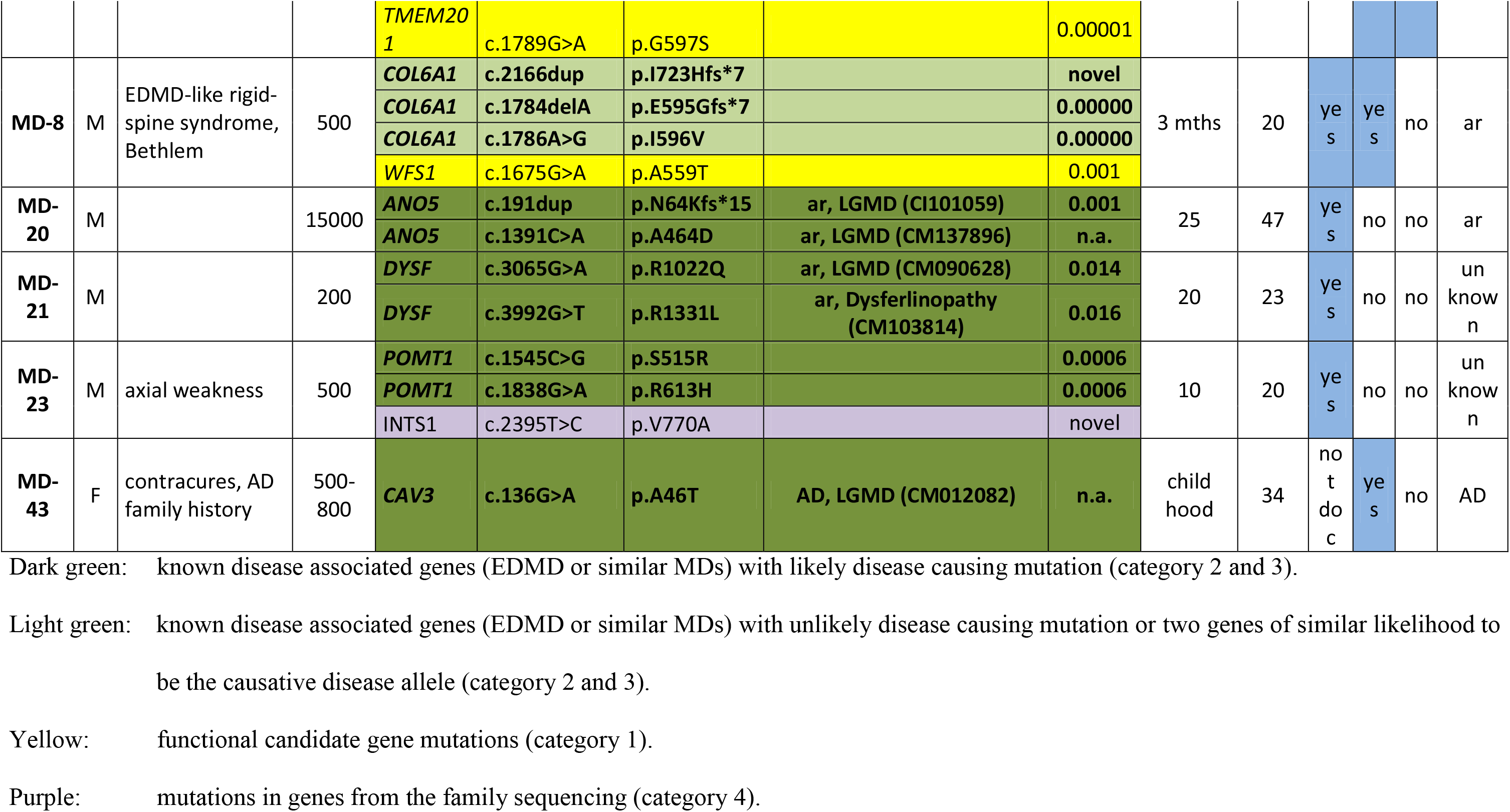
Solved Patients

Gene category II of related muscular dystrophies yielded 18 patients with mutations considered causative. Four of these patients had combinations of a missense and frameshift mutation in *CAPN3* (Table 1). *GBE1* mutations were found in four other patients: three missense and one splice-site. *VCP* and likely recessive *TTN* were mutated in two patients each; however, *TTN* mutation patients also carried *SYNE1* mutations. Genes with one patient carrying likely disease-causing mutations were *COL6A1*, *CAV3*, *DMD*, *ANO5*, *DYSF* and *POMT1*. The *DMD* mutation created a stop codon at codon three, resulting in possible usage of an alternative start codon and a milder phenotype than Duchenne^26^. For *ANO5*, *DYSF* and *POMT1* the respective patients had two mutations, consistent with the reported inheritance (autosomal recessive for MD-20/*ANO5* and unknown for MD-21/*DYSF* and MD-23/*POMT1*; Table 1). However, lacking DNA from the parents we could not perform segregation studies.

Several category III genes from exome sequencing were elevated to strong candidates if mutated in multiple patients within the primer library cohort based on the assumption that causative genes will be independently mutated in multiple patients. The top candidates were *INTS1*, *ANK2*, *XIRP1* and *USP34*. Heterozygous *ANK2* mutations were identified in family 5 plus six cohort patients with no other obvious disease-causing mutations and so were most likely causative (Table 2). Causation is similarly likely for other genes; however, in some patients there were additional candidate alleles identified. Heterozygous *INTS1* was mutated in four members of family 3 plus five cohort patients, four of whom had no mutation in already associated genes (Table 2). The last patient, MD-23, additionally carried two *POMT1* mutations; however, it is unclear if the likely recessive *POMT1* mutations affected one allele or both so causation remains undetermined. Other good category IV candidates were *USP34* (heterozygous mutations in exome sequenced family 2) and *XIRP1* (mutated in families 2 and 4), each with mutations in an additional five patients. Some patients had additional mutations in already associated genes, but if these other mutations were causative then modifying effects for the new candidates are still possible.

**Table 2:**
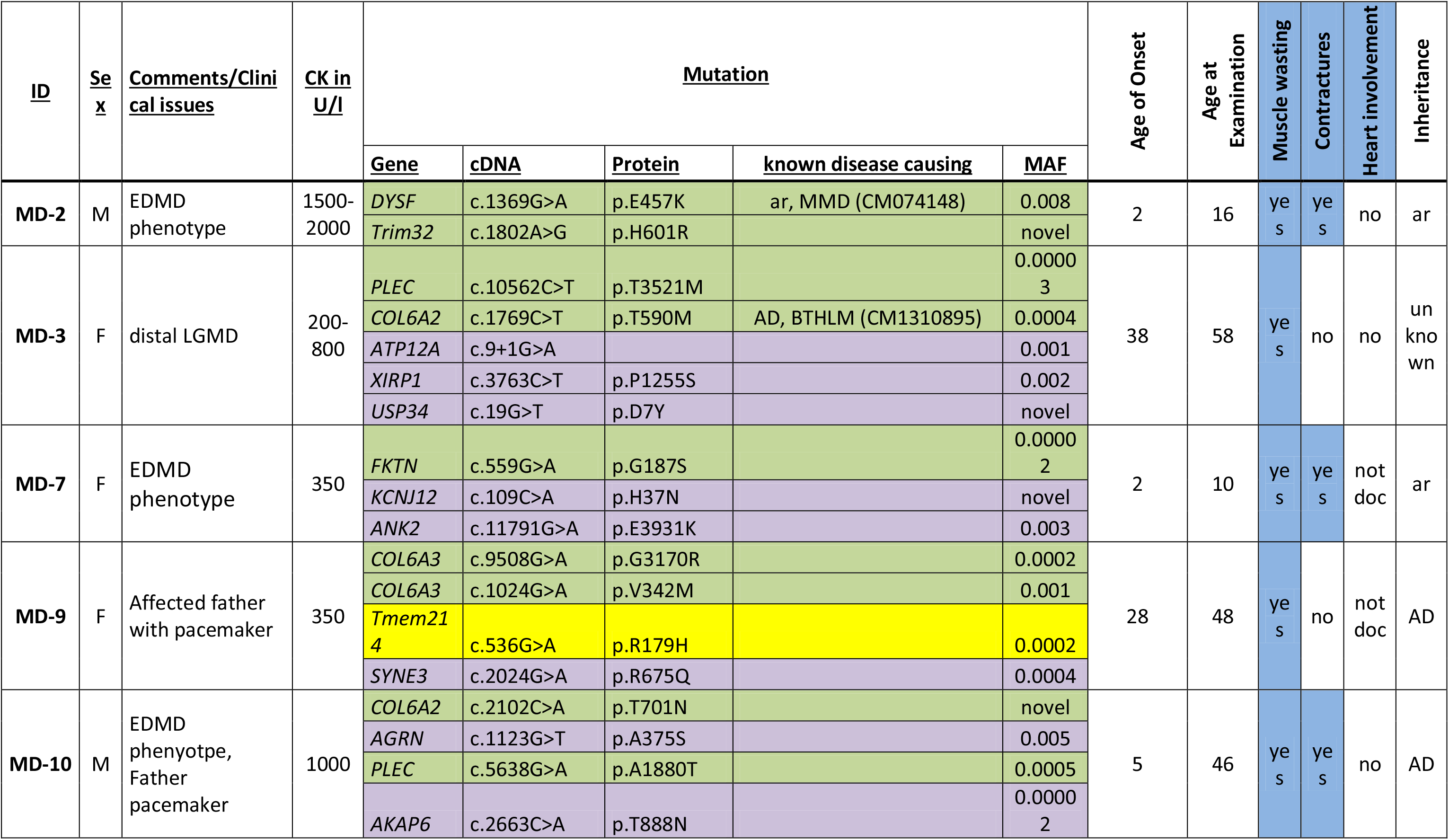
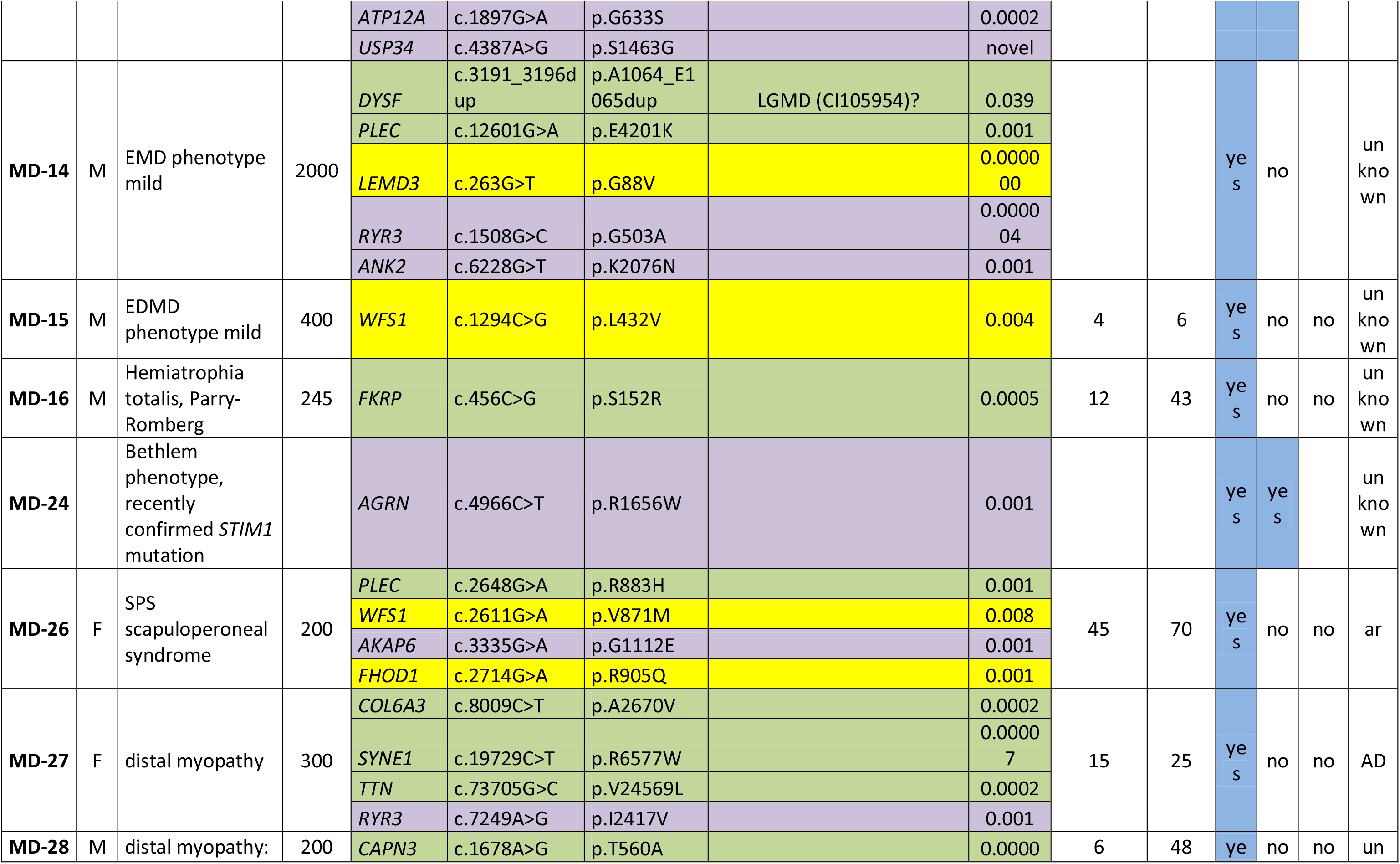
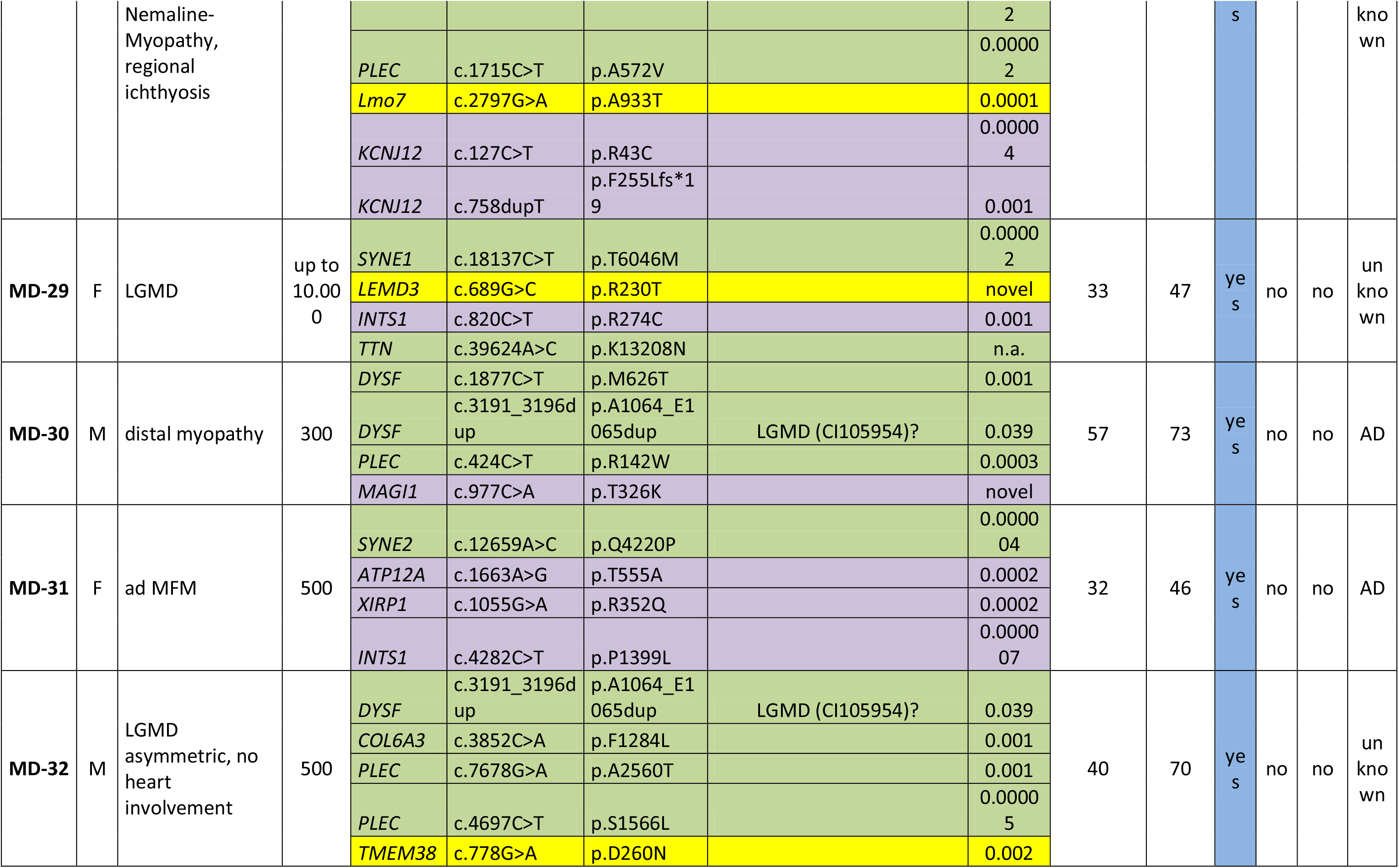
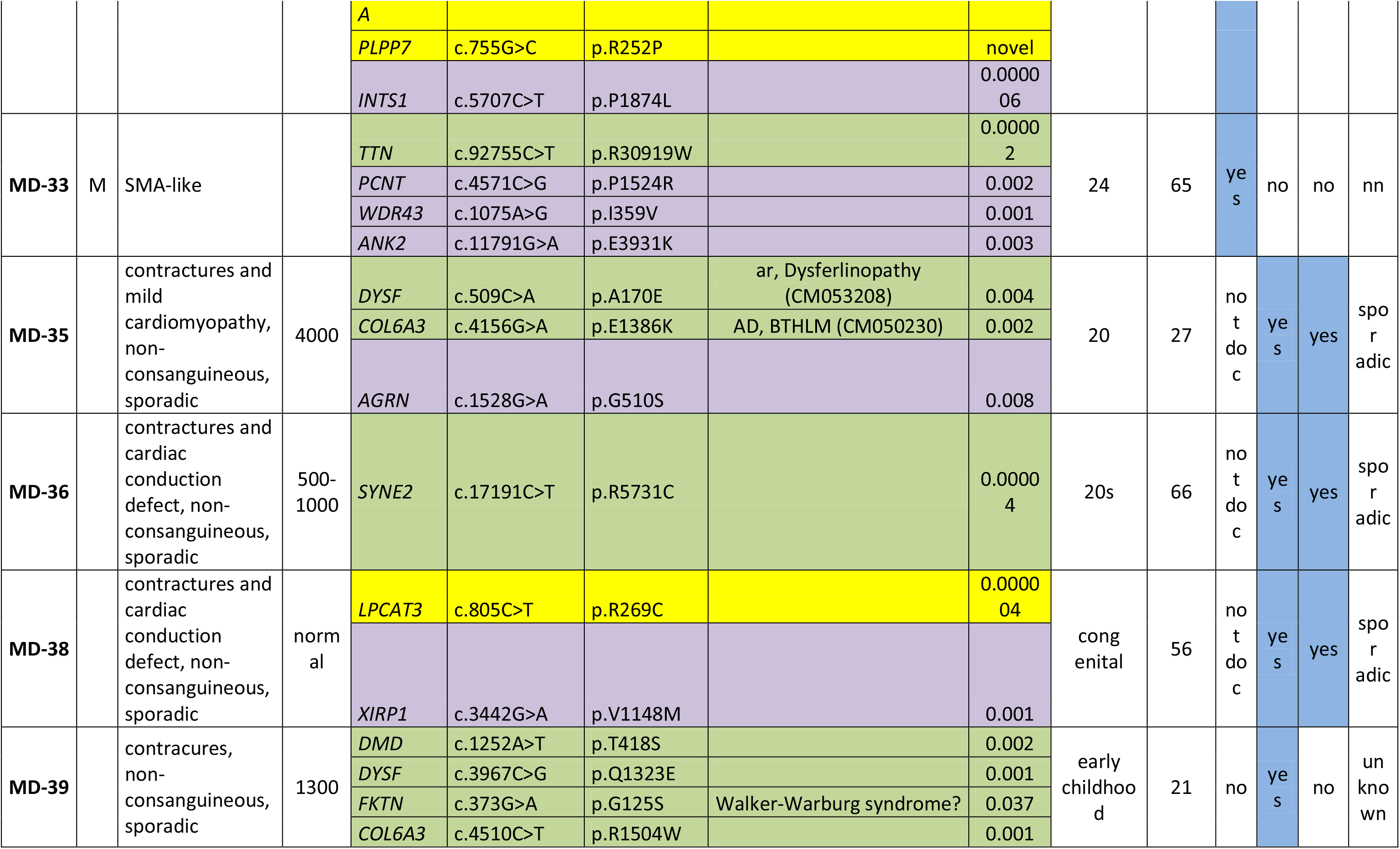
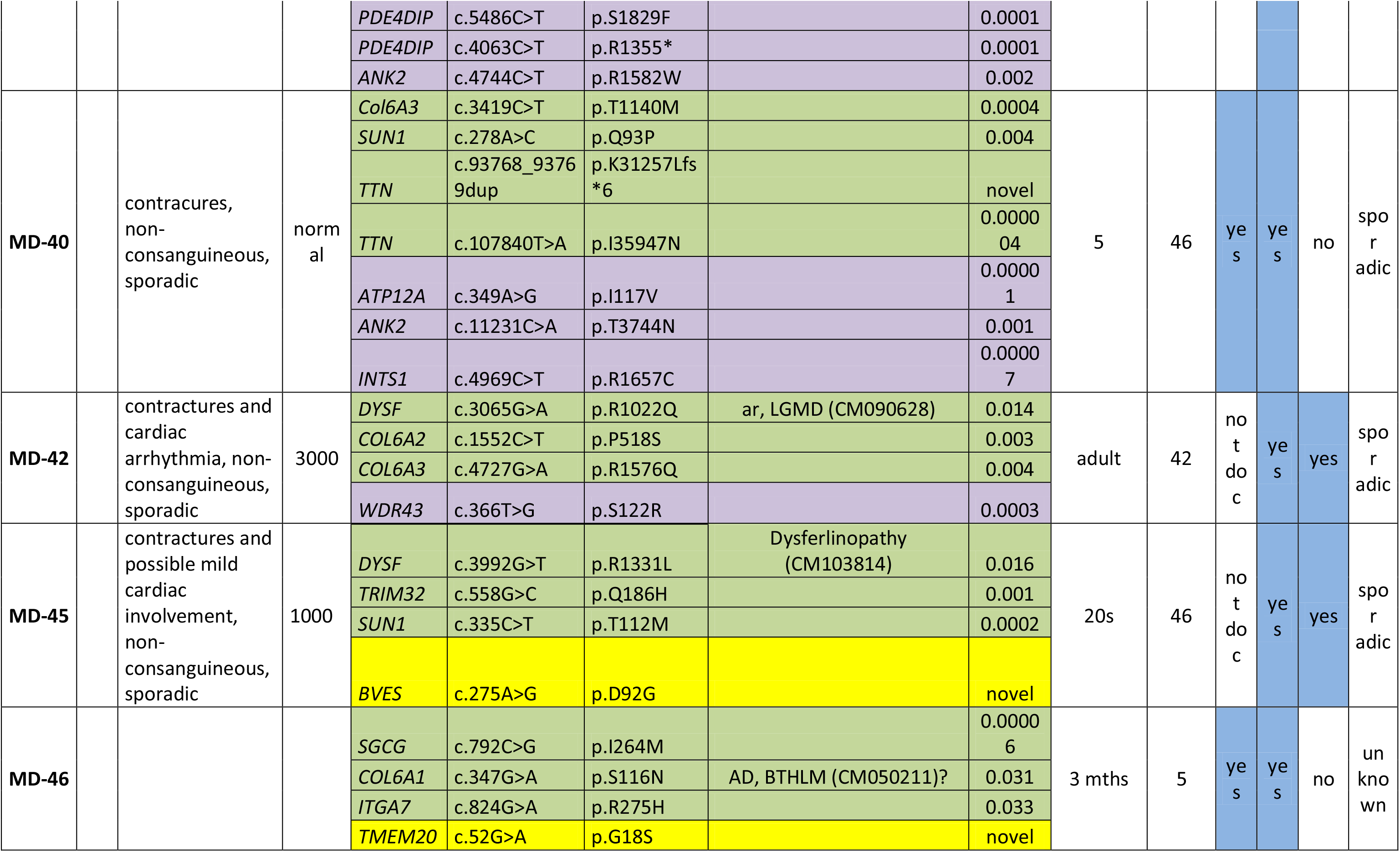
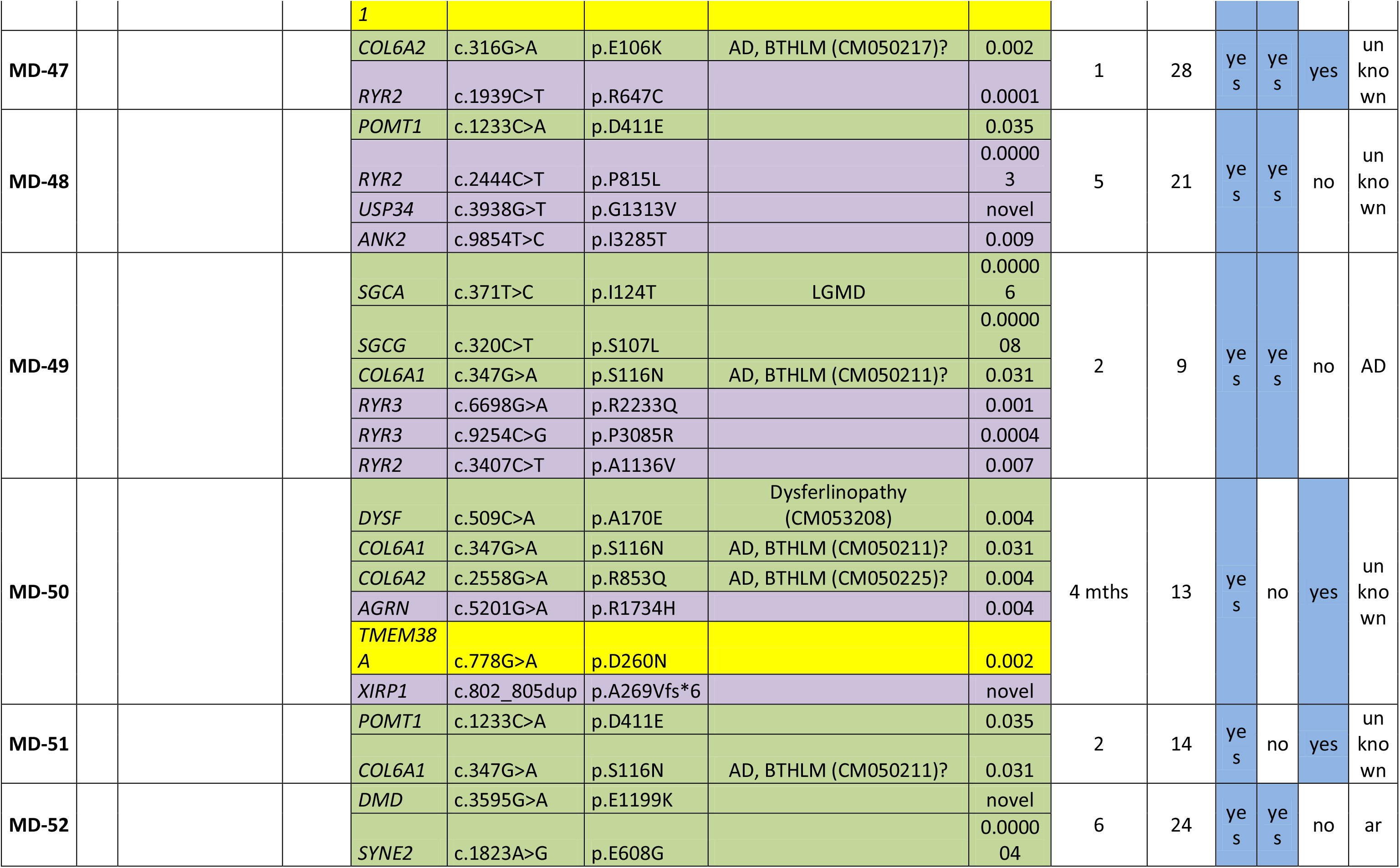
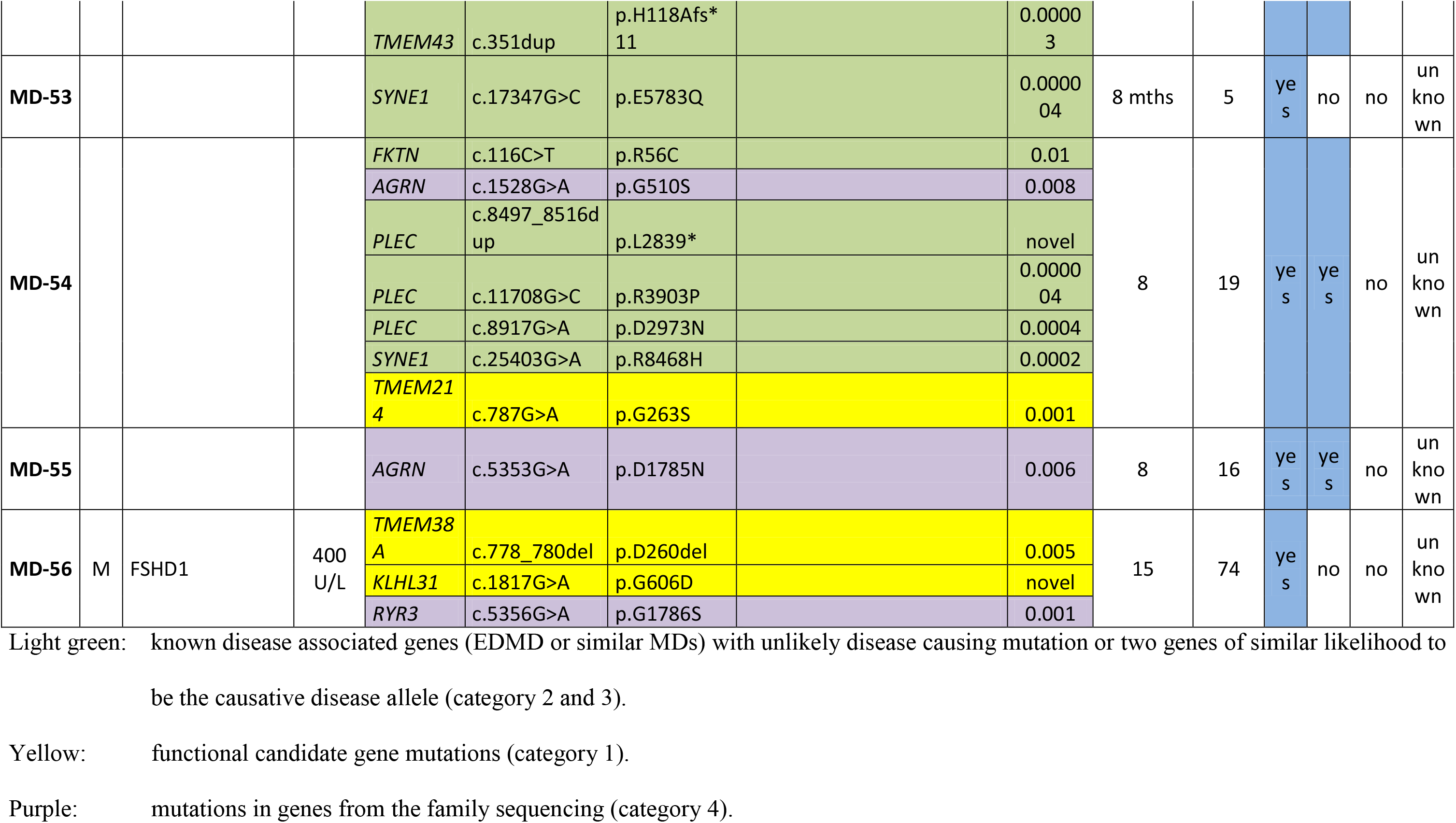
Patients with Candidate Genes

Several category IV functional/tissue candidate genes were mutated in 16 of the 56 primer library cohort patients. These were *WFS1* (4 patients), *TMEM201* (3 patients), *TMEM38A* (3 patients), *PLPP7* (2 patients), *TMEM214* (2 patients), *LPCAT3* (1 patient), *KLHL31* (1 patient), and *BVES* (1 patient). Of these, three patients with *TMEM38A* mutations, two patients with *TMEM214* mutations, one patient with an *LPCAT3* mutation and one patient with a *BVES* mutation were clearly the top candidates with no other reasonable candidates identified and patient MD-32 carried mutations in both *TMEM38A* and *PLPP7*. Other mutations identified were in association with other possible candidates that included likely causative mutations in *GBE1*, *COL6A1*, *LMNA* and *TTN* (details in Table 1). The patient with the combined *LMNA* and *TMEM201* mutations had a very early age of onset (1 year), suggesting that both mutations contribute to the more severe (congenital) phenotype as the *LMNA* mutation has not been associated with congenital muscular dystrophy.

All in all, sequencing the 56 additional patients with the primer library found mutations in only a subset of the 252 candidates from the exome sequencing and this subset is expected to be much higher confidence because causative genes are more likely to be also mutated in other EDMD patients. In contrast, mutations were found in 19 of 25 related muscular dystrophies and in 11 of 16 functional candidates; so a strong enrichment for these candidate pools was observed (Fig. 2A).

### NUCLEAR ENVELOPE LINKS

All previously linked EDMD genes encode nuclear envelope proteins. The functional candidates were also biased towards genes identified in the nuclear envelope by proteomics; however, there was no bias towards the nuclear envelope when selecting genes for the primer library from similar muscular dystrophies or from exome sequencing. Nonetheless, the majority of genes from similar muscular dystrophies encode proteins for which at least a subpopulation associates with the nuclear envelope (Fig. 2B). Interestingly, just considering the candidates from the exome sequencing in which mutations were also found in other patients from the primer library sequencing, the nuclear envelope portion increased from less than 10% to more than 40% - considerable more than the overall genome portion of 5.9% (Fig. 2B). Of note, the proteins encoded by genes linked to other muscular dystrophies such as *COL6A1*, *CAV3*, *DYSF*, *DMD*, *TTN*, and *VCP* and the strongest family sequencing candidates *INTS1* and *ANK2* were all found in nuclear envelope proteomics datasets^20, 27^. While these could reflect either a separate pool in the nuclear envelope or connections that were maintained during nuclear envelope isolation, this suggests at least an indirect physical connection of these candidates to the nuclear envelope.

The two top argued mechanisms for how mutations in nuclear envelope proteins can cause pathology are mechanical instability and genome misregulation. Genes in different candidate categories contained Gene Ontology (GO)-terms for functions in gene regulation, cytoskeleton, and both together. Interestingly, the likely candidates from all categories were enriched for genes simultaneously linked to both gene regulation and cytoskeleton GO-terms compared to the overall genome (Fig. 2C). Such genes may be involved in mechanosignal transduction to the genome. Consistent with this idea, most of the proteins encoded by the final candidate genes interact with other candidates according to interactome studies and these interactions form a chain of connectivity between the nuclear envelope and the plasma membrane via cytoskeletal proteins that could support mechanotransduction to the genome (Fig. 2D).

### CONFIRMATION OF NOVEL EDMD ALLELES

Thus far only the three *LMNA* mutations, the *CAV3* and one of the *CAPN3* (MD-43) mutations have been confirmed as insufficient numbers of family members have come to clinic for linkage analysis. Therefore, to test the likelihood that other mutations identified cause EDMD disease pathology, we tested two of the gene regulating NETs to determine if the mutations identified disrupt their normal functions in myogenic gene regulation. In keeping with this idea, for the 8 out of 16 functional NET candidates where mutations were found (6 of which have known gene regulation functions), nearly all mutations identified faced the nucleoplasm or were positioned where they could alter membrane topology (Fig. 3A). The two muscle-specific NETs we chose to test were PLPP7/NET39 and Tmem38A. Both recruit partially overlapping, but mostly different sets of genes to the nuclear periphery to enhance their repression and many of the genes targeted are antagonistic to myogenesis or from alternate differentiation pathways^24^. Combined knockdown of PLPP7/NET39, TMEM38A and WFS1 blocked myogenesis, providing a logical route from their disruption to muscle disease pathology. Therefore, the *PLPP7* and *TMEM38A* mutations were exogenously expressed in C2C12 myoblast cells to determine if they could perform the previously shown gene positioning function of the wild-type in recruiting specific gene targets to the nuclear periphery for enhanced repression. Tmem38A normally repositions the *DDR2* gene locus to the nuclear envelope to repress it during myogenesis, but with mutations p.N260D and p.N260del it fails to do so (Fig. 3B). Similarly, PLPP7/NET39 normally recruits the *PTN* gene locus to the nuclear envelope to repress it during myogenesis, but with mutation p.R252P it could not. PLPP7/NET39 mutation p.M92K also affected the gene positioning, though apparently in the opposite direction which might also affect expression (Fig. 3C).

**Figure 3.**
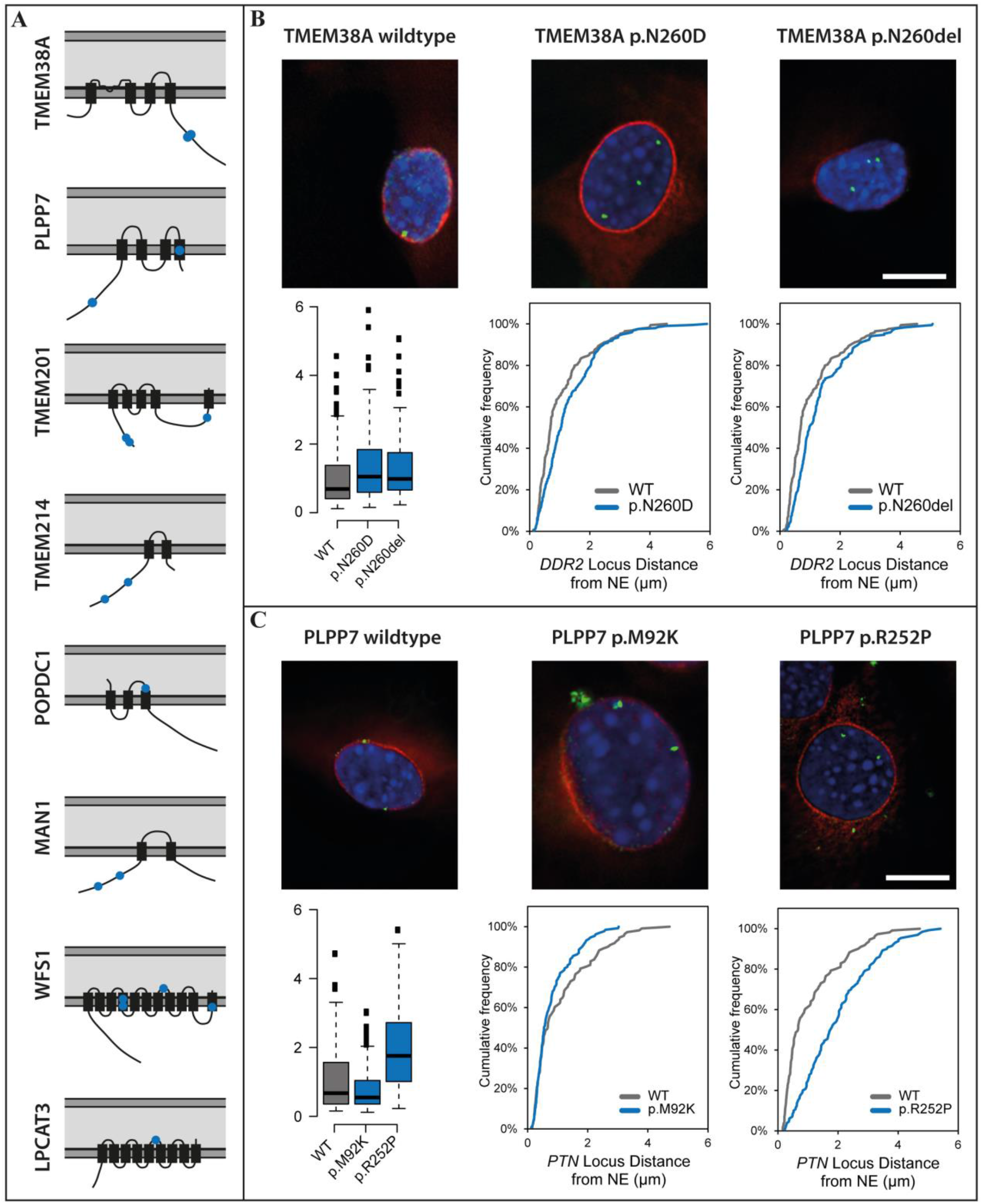
Mutations in muscle gene-repositioning NETs affect their ability to recruit genes to the nuclear envelope (NE). **(A)** Schematic presentation of the topology of further muscle NETs and their mutations identified by the primer library sequencing. The lipid bilayers of the nuclear envelope are shown in dark grey and the lumen of the nuclear envelope in light grey. Transmembrane segments are thicker black rectangles and point mutations identified are shown in blue. The mutations identified are all positioned in nucleoplasmic regions where they could either interact with the genome or at transmembrane spans where they could disrupt protein topology and hence also genome interactions. **(B)** FISH showing the localization of the *DDR2* gene (green) in C2C12 mouse myoblasts upon the expression of RFP-tagged wild type and mutant TMEM38A that can be seen in both cases to target to the nuclear envelope (red, upper panel). The cumulative frequency of the distance of the gene loci to the NE for each mutation compared to the wild type is shown under each image of the cells expressing the mutant NETs and a whisker plot summary for the distance to the NE of all mutations is given in the lower left corner. Both mutations block the ability of TMEM38A to reposition the *DDR2* locus to the NE. **(C)** FISH showing the localization of the *PTN* gene (green) in C2C12 mouse myoblasts upon the expression of GFP-tagged wild type and mutant PLPP7 (red, upper panel). Cumulative frequency plots of the distance of the gene loci to the NE for each mutation and the summary for the distance to the NE of all mutations are given as in *B*. The mutations also affect the gene repositioning function of PLPP7.

## Discussion

Failure of high throughput genomic approaches to identify new disease alleles can at least in some cases be overcome by our multistage approach. This approach pinpointed candidates in part based on the preferential tissue focus of pathology and in part on the subcellular localization of known alleles. Similarly applying filters in focusing candidates for such a multipronged approach can be applied to other genetically heterogeneous diseases.

With nearly half of EDMD cases previously linked to genes encoding 6 nuclear envelope proteins it was clear that EDMD is a nuclear envelope disease. This is strengthened by enrichment for nuclear envelope proteins amongst our top new candidate alleles. *COL6A1*, *CAV3*, *DYSF*, *DMD*, *TTN*, and *VCP* gene products were found in muscle nuclear envelopes^20^. As most of these proteins have previously been associated with the cytoskeleton or plasma membrane, their association with the nuclear envelope may be indirect through lamin-cytoskeletal connections. However, this association could also be due to splice variants that target to the nuclear envelope or specific translocation to the nucleus under certain conditions as has been shown for *CAV3* family member caveolin 2. In this case, a caveolin 2 subpopulation translocates to the nucleus and interacts with lamin A to regulate histone modifications and gene expression^28^.

The gene positioning defects for *TMEM38A* and *PLPP7* mutations not only further link the nuclear envelope to EDMD, but also strengthen the idea that misregulation of myogenic gene expression is the primary cause of EDMD pathology. In addition to Tmem38A and Plpp7, the muscle NETs Tmem214, WFS1, and NET5/Samp1 all have gene-repositioning functions that contribute to gene regulation and the NET MAN1 affects gene regulation through its interactions with Smads as well as binding several chromatin partners^29, 30^. The involvement of these muscle gene repositioning NETs, not only as novel causative alleles but also in mediating EDMD pathology caused by mutations in widely expressed nuclear envelope proteins, is further supported by WFS1, Tmem214, Tmem38A, and NET5/Samp1 being mislocalized in isolated differentiating EDMD muscle cells or muscle biopsy sections^31^.

Of the previously EDMD-linked nuclear envelope proteins, Lamin A has both cytoskeletal and genome regulation functions; so its mutation could support both mechanical instability and genome misregulation hypotheses for EDMD pathophysiology^32–37^. Emerin interacts with actin supporting a cytoskeletal role, but it also has many reported contributions to genome regulation through its binding DNA condensing factors BAF and HDAC3, splicing factors, the transcription factor Lmo7, and the transcriptional repressor germ cell-less^38^. FLH1 is linked to signal transduction and splice variant FHL1B targets specifically to the nuclear envelope^39^. Moreover, FHL1 has been linked to other myopathies such as X-linked myopathy with postural muscle atrophy (XMPMA)^40^ via its signal transduction function.

As signaling functions could affect both gene regulation and the cytoskeleton, these mechanisms toward pathology were considered equally likely; however, a gene mis-regulation mechanism is much more likely now with the new gene-repositioning candidate alleles identified. Though there are some other disparate functions reported for several of these NETs^20, 22, 41^, WFS1, Tmem38A/TRIC-A, NET39/ PLPP7, Tmem214 and NET5/Samp1 are all at the nuclear envelope preferentially in muscle and all share a common function in directing gene-repositioning for regulation of gene expression during myogenesis^24^. That some of these muscle-specific NETs had overlap in their functions further supports the possibility of their working in a common pathway towards EDMD pathophysiology. At the same time, while there was some overlap in the sets of genes regulated by these muscle NETs, each had also unique gene targets. The links of candidate alleles to gene repositioning and mechanotransduction are the more compelling in this context because the different sets of genes regulated — all important in myogenesis — thus supports the clinical variation observed in EDMD.

Our sequencing in patients diagnosed with an EDMD-like phenotype identified mutations in several genes linked to muscular dystrophies that share clinical features with EDMD. This might reflect incorrect diagnoses or their involvement in EDMD. The latter case seems likely, considering that *COL6A1*, *CAV3*, *DYSF*, *DMD*, *TTN*, and *VCP* gene products link to the nuclear envelope. Indeed, many of these gene products interact with one another in a way that could form a chain from the plasma membrane to the nuclear envelope (Fig. 2D). This also is compelling to this gene regulation mechanism as this chain could play a role in mechanosignal transduction to the nucleus.

Finally, as the families chosen for exome sequencing all had differences in presentation, there are likely additional mutations picked up in the primer library that eventually might be used as predictors of severity or other aspects of clinical presentation once further sequencing reveals better correlations. Thus it would be beneficial to continue using this primer library diagnostically both to find these correlations and because it is cheaper and faster than standard iterative Sanger sequencing for such a genetically variable disease to identify mutations in known linked genes. In general, this iterative multipronged approach, combining into a primer library a set of preliminary candidates from exome sequencing in which only sufficient pedigrees exist for partial linkage analaysis together with candidates from related disorders and candidates specific to the tissue where pathology is manifested that are associated with linked organelles and functions, might be applied to a wider range of genetically heterogeneous orphan diseases where insufficient numbers of patients are available for standard genome and exome approaches to be effective.

## Materials and Methods

### PATIENT MATERIALS AND ETHICS

Patient DNA was obtained from the Muscle Tissue Culture Collection (MTCC) at the Friedrich-Baur-Institut (Department of Neurology, Ludwig-Maximilians-University, Munich, Germany), the Institute of Human Genetics, University of Newcastle upon Tyne, Newcastle upon Tyne, UK, the MRC Centre for Neuromuscular Disorders Biobank (CNDB) in London, the Department of Pediatric Neurology, Developmental Neurology and Social Pediatrics at the University of Essen, the Rare Diseases biological samples biobank at the Dubowitz Neuromuscular Centre, Great Ormond Street Hospital for Children NHS Trust, London, UK. All materials were obtained with written informed consent of the donor at the CIND, the CNDB or the MTCC. Ethical approval for the Newcastle MRC Centre Biobank for Neuromuscular Diseases is covered by REC reference 08/H0906/28+5 and IRAS ID 118436 and MTA CT-2166, that of the Rare Diseases biological samples biobank for research to facilitate pharmacological, gene and cell therapy trials in neuromuscular disorders is covered by REC reference 06/Q0406/33 with MTA reference CNMDBBL63 CT-2925/CT-1402, and for this particular study was obtained from the West of Scotland Research Ethics Service (WoSRES) with REC reference 15/WS/0069 and IRAS project ID 177946.

### EXOME, RNA, AND GENOME SEQUENCING

Genome: 15X clean depth coverage using 90PE Illumina HiSeq2000 technology. RNA-Seq: total RNA from biopsy tissue with rRNA depletion and random-primed cDNA preparation and PE100 sequencing on a Hi-Seq2000 platform with 20 million reads minimum (Otogenetics Corporation, Norcoss, USA).

Exome: Sequencing was performed on the Illumina HiSeq and raw data processed with CASAVA 1.8.

### FLUORESCENCE *IN SITU* HYBRIDIZATION

Mutations were generated by Agilent Site-Directed mutagenesis. Plasmids encoding tagged Tmem38a, PLPP7 and mutants were transfected using Lipofectamine 3000 (Invitrogen) into C2C12 cells (ATCC, VA, USA) cultured at 37°C, 5% CO2 in DMEM containing 20% FBS, 50U/mL penicillin and 10mg/mL streptomycin. Fluorescent *in situ* hybridization (FISH) experiments were performed as described in^42^.

### PRIMER LIBRARY CONSTRUCTION, PROCESSING AND SEQUENCING

A SureSelect^XT^ Custom 1.638 Mbp target enrichment library (5190-4817) containing 25,036 oligonucleotide probes against H. sapiens hg19 GRCh37 sequence as of February 2009 was prepared by Agilent for use with Illumina multiplexed sequencing platforms. Patient genomic DNA was isolated from blood and prepared for sequencing using the SureSelect^QXT^ Reagent Kit (G9681B) according to the manufacturer’s instructions. Recommended minimum sequencing per sample was 327.793 Mbp and an average of 3,427,092 was obtained with a range from 442,125 to 7,066,507 using 125 base paired-end sequencing on a Hi-Seq2500.

### BIOINFORMATICS

Variant analysis was performed using the Genome Analysis toolkit [GATK] v2.7-2^43^ and picard tools v1.74 (http://broadinstitute.github.io/picard/) using GATK Best Practices recommendations^44, 45^ against human genome assembly hg19. The allele frequencies of variants were cross-referenced with gnomAD version 2.1^46^ using both the genome and exome datasets.

RNA-Seq: STAR v2.1.1^47^ was used to map reads to the hg19 reference genome, samtools v0.1.19^48^ was used for file conversion. Deeptools v1.5.1^49^ was used for downstream analysis.

## Acknowledgements

We dedicate this study in honor of Alan E.H. Emery who not only described X-linked EDMD as a distinct disease in 1966 but also initiated the longest standing regular Laminopathy meeting in 2006. We further want to thank all patients and their families for their valuable contribution. Funding for this work was principally provided by Wellcome Trust Grants 095209, Muscular Dystrophy UK grant 18GRO-PG24-0248, and MRC grant MR/R018073/1 to ECS and 092076 for the Centre for Cell Biology. Funding was also provided by the European Community’s Seventh Framework Programme (FP7/2007-2013) “Integrated European –omics research project for diagnosis and therapy in rare neuromuscular and neurodegenerative diseases (NEUROMICS)” (grant agreement n° 2012-305121); the Muscular Dystrophy UK Grant on Gene Identification to FM, and the support of the MRC Neuromuscular Centre Biobank at UCL is also gratefully acknowledged.

## Author contributions

ECS, MW and PM designed the project. PM performed the sequencing and data analyses. RC performed the FISH experiments. ARWK and JIH performed the bioinformatics. EH, HK, FM, US, VS and BS contributed patient material and clinical description. PM and ECS wrote the paper. All authors read the manuscript, offered feedback, and approved it before submission.

## Supplementary Material

Table S1: Family Exome Sequencing Results

Table S2: Genome Sequencing Index Patient Family 1 Results

Table S3: RNA Sequencing Index Patient Family 1 Results

Table S4: List of Genes in Primer Library

Table S5: Primer Library Sequencing Results

